# Scale invariance of BMP signaling gradients in zebrafish

**DOI:** 10.1101/489872

**Authors:** Yan Huang, David Umulis

## Abstract

In both vertebrates and invertebrates, spatial patterning along the Dorsal-ventral (DV) embryonic axis depends on a morphogen gradient of Bone Morphogenetic Protein signaling. Scale invariance of DV patterning by BMPs has been found in both vertebrates and invertebrates, however the mechanisms that regulate gradient scaling remain controversial. To obtain quantitative data that can be used to address core questions of scaling, we introduce a method to tune the size of zebrafish embryos by reducing varying amounts of vegetal yolk. We quantified the BMP signaling gradient in wild-type and perturbed embryos and found that the system scales for reductions in cross-sectional perimeter of up to 30%. Furthermore, we found that the degree of scaling for intraspecies scaling within zebrafish is greater than that between Danioninae species.

## Introduction

An important area in developmental biology focuses on identifying mechanisms of robustness to perturbations that organisms face during development. One key variable that persists in many systems is the ability of the patterns to form in proportion to the size of the domain in which they are active. The ability of the system to adapt to different size domains has also gained a lot of interest from physicists and engineers because the biophysical processes thought to be responsible for developmental pattern formation do not automatically account for differences in size and would instead predict a pattern that would not scale. Recent advances have introduced new hypotheses regarding scaling, however in many systems the mechanisms of scaling remain poorly understood [1, 2, 3]. In invertebrates, quantitative evidence of scale invariance has been reported in Drosophila in multiple instances. Dynamic scaling of the Decapentaplegic (Dpp) gradient is reported during growth of the wing imaginal disc in *Drosophila melanogaster* [4]. In flies, both interspecies and intraspecies scaling occurs for the anterior-posterior embryonic patterning by Bicoid (Bcd) and for dorsal/ventral embryonic patterning by Dpp [5]. Interspecies scaling is observed in patterns of gap and pair-rule gene expression in blastoderm embryos of *Lucilia sericata, Drosophila melanogaster*, and *Drosophila busckii* by Bicoid (Bcd) protein [6]. Intraspecies scaling of tissue development along the anterior-posterior axis in *Drosophila melanogaster* is reported to be traced back to the scaling of anterior Bcd production rate with embryo volume [7]. These examples of scaling have also provided new insight on mathematical modeling of pattern formation and scaling [5, 8].

In vertebrates, the phenomenon of intraspecies scaling has been reported in skin pattern formation in fish pigmentation, and under bisection in Xenopus and goldfish embryo development but the gradients in these systems have not been quantitatively measured to identify whether the gradients themselves scale or if the downstream correction contributes to the observed scaling [9,10, 11, 12]. Interspecies scaling in neural tube patterning between the zebra finch and the chick is suggested as being traced back to the ratio of activating and repressive transcription factors of the morphogen Sonic Hedgehog (SHH) [13].

Our focus herein is to quantify dorsal/ventral scaling by measuring signaling gradients in zebrafish embryos. Embryonic DV patterning is mediated by the extracellular distribution of Bone Morphogenetic Proteins (BMPs). The BMP morphogen gradients are established by a network of extracellular regulators including antagonists that bind and inhibit ligand-receptor interactions and regulate the formation of the gradient in both vertebrates and invertebrates [14, 15]. BMP gradient formation has been studied and several potential mechanisms for how the antagonists shape the gradients have been proposed in Drosophila [16, 17, 18, 19, 20, 21, 22], Xenopus [13, 23, 24], zebrafish [16, 25, 26] and mouse [27].

To determine whether DV patterning by BMPs in zebrafish is scale invariant, we developed a protocol to physically reduce the size of embryos. We measured the BMP signaling gradient using quantitative imaging and we measured the degree of scaling in zebrafish DV patterning. This not only allows us to identify the scale invariance phenomenon in zebrafish embryonic DV patterning but also opens a window for us to further investigate the formation and regulation of BMP gradients by perturbing size. In addition to surgical size modifications of sibling *Danio rerio*, we quantified morphogen gradients and the degree of scaling of DV patterning between zebrafish (*Danio rerio*) and another closely related species giant danio (*Devario aequipinnatus*, abbreviated by Gd) in the Danioninae clade. Through experiments and quantitative analysis, we found that zebrafish embryos are able to scale for a range of size reductions, however scaling does not persist between Gd and zebrafish. We provide a new paradigm for discovering scale invariant systems and investigating potential mechanisms, as well as new metrics to quantify the degree of scale invariance and compare scaling among different systems.

## Results and discussion

### Intraspecies scaling of zebrafish embryonic DV patterning

We generated Cut embryos by removing vegetal yolk between the 8-cell and 256-cell cleavage stages, and split the eggs into two groups, one group of eggs was fixed at shield stage for staining, the other group was placed in the incubator to observe the phenotypes at 1 dpf. From this experiment, we find a distribution of phenotypes including many that are smaller than WT that display a proportional DV axis the same as WT. Figure 1a plots the frequency of Cut embryos and displays different phenotypes after yolk removal surgery. Over 30% of the Cut embryos that are 30~50% smaller than WT display a normal phenotype, and the ratio is higher than 50% for Cut embryos that have smaller perturbations ranging from 0~30% smaller than WT. The distribution of embryos suggest that the system is capable of scaling, however the level of survival decreases as the perturbation increases.

Figure 1b shows the normal phenotype and other phenotypes among Cut embryos at 1 dpf (Right) and their corresponding images at 6 hpf (Left). The Cut embryos are fixed at the same time when WT grows to shield stage (6 hpf). Figure 1b displays normal, C1, C2, C3, C4, and C5 phenotypes among the Cut embryos. The Cut embryo that displays a normal phenotype at 1 dpf is about 13% smaller in length and 33% smaller in volume than WT embryo at 6 hpf. This is an example of pattern scale invariance among zebrafish embryos of different sizes. The last embryo in Figure 1b shows a Cut larva with a ventralized phenotype and no head.

**Figure 1.**
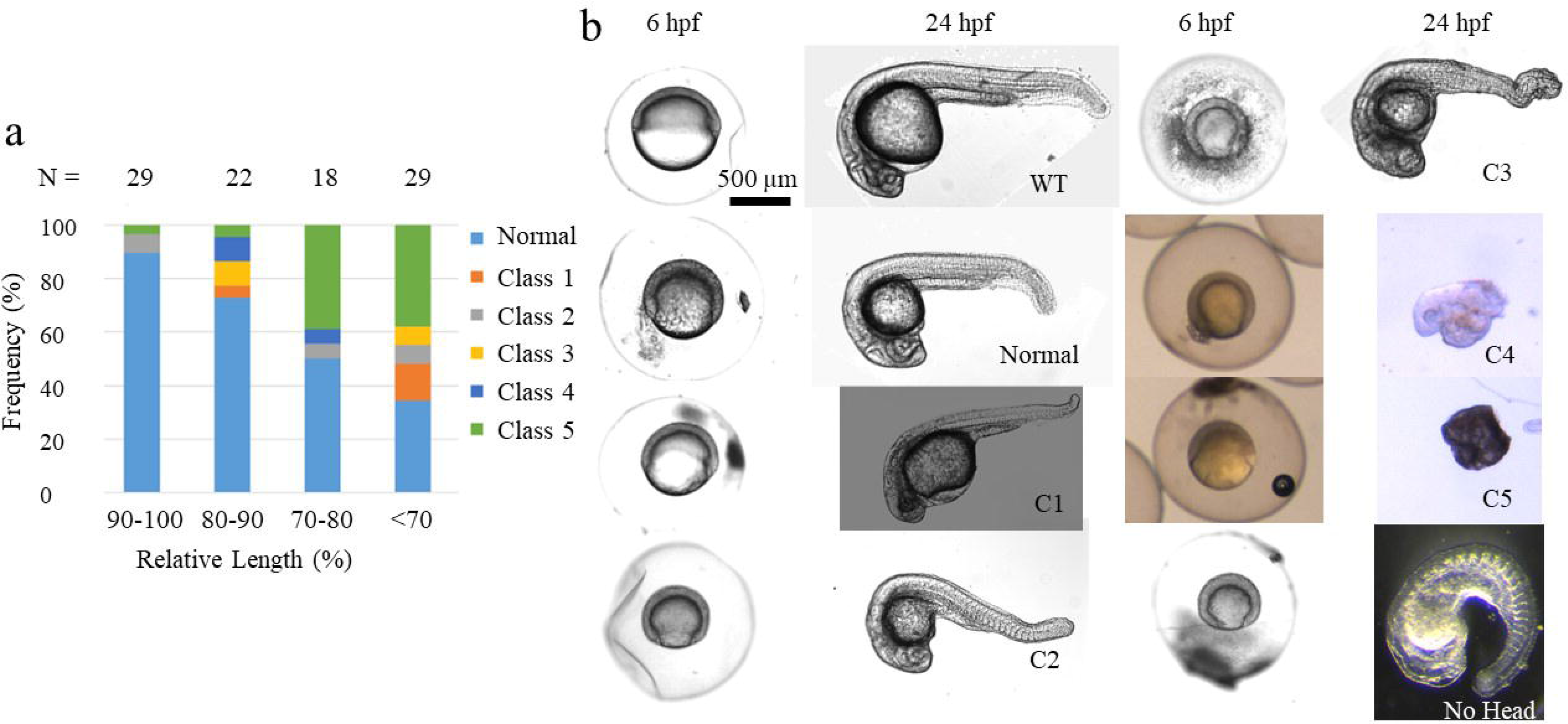
Phenotype distributions of cut embryos. **(a)** The bar plot shows the frequencies of different phenotypes of Cut larvae at 1 dpf. Normal: DV proportions same as wild-type. Class 1~5: dorsalized phenotypes from weakest to strongest, with the proportions of DV axis tissues different from WT or some tissues along DV axis missing. The definitions of these 5 classes of phenotypes are used as in Mullins et al. Class 1: no tail fin. Class 2: no tail vein. Class 3: no yolk extension. Class 4: no tail. Class 5: the yolk spills out of the embryo and the embryo lyses [33]. The number of embryos that look ventralized is also combined to Class 5 category. **(b)** Lateral view of live WT (top left two) and Cut embryos at 6 hpf and 24 hpf under bright-field microscope. Embryos displaying normal, C1, C2, C3, C4, C5(the embryo lyses) and ventralized (no head) phenotypes are imaged at 6 hpf and 24 hpf with phenotype labels on the 24 hpf image panel.

To determine if scaling of the dorsal/ventral gradient of BMP signaling is achieved in early development, we quantified the Psmad gradient in populations of cut embryos. Figure 2a shows the result after segmentation that yields an embryo point cloud with Psmad labeling intensity shown by the color heatmap for WT and two Cut embryos. The middle Cut embryo displays a similar Psmad gradient as the WT embryo while the ventral Psmad labeling intensity of the rightmost Cut embryo in Figure 2a has much lower intensity than that of the WT embryo.

**Figure 2.**
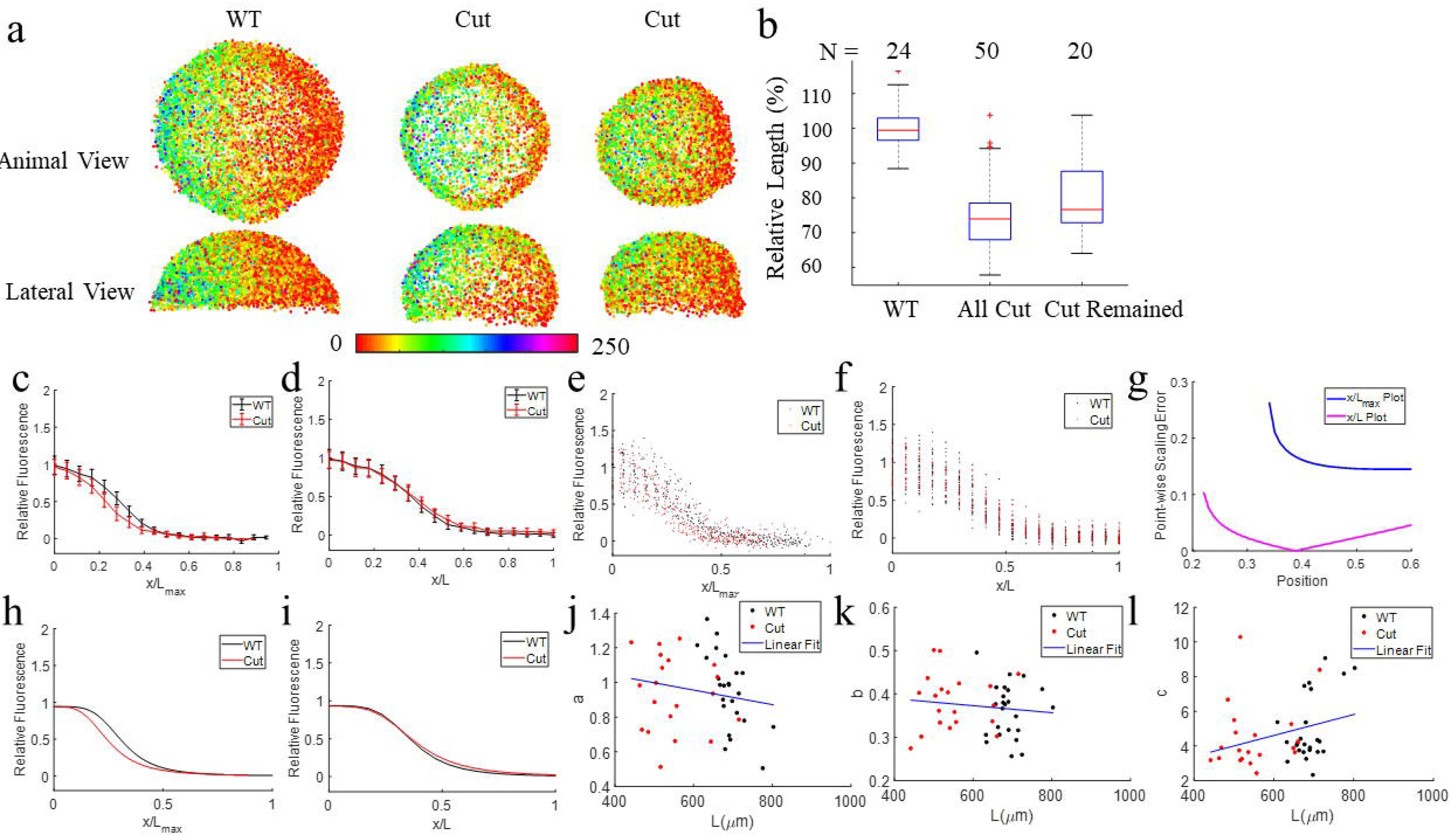
Psmad gradient profiles of WT and Cut scale. **(a)** The animal and lateral views of Psmad stained 6 hpf embryo point clouds after image processing with Psmad labeling intensity in color. The animal and lateral view of a Psmad stained WT embryo (Left) displays a similar Psmad gradient as the Cut embryo (Middle). The animal and lateral view of a Psmad stained Cut embryo (Right). **(b)** The boxplot of relative egg sizes of Psmad stained embryos. From left to right, they are all WT (N = 24), all Cut (N = 50) and the Cut embryos that pass the filtration rules (N = 20). **(c)** – **(f)**: Psmad gradient profiles of WT (black) and Cut (red) embryos after filtration. **(c)** Population mean Psmad gradients of WT and Cut after filtering on x/L_max_ plot with variation shown by error bar. **(d)** Population mean Psmad gradients of WT and Cut on x/L plot with variation shown by error bar. **(e)** Psmad signal of embryos grouped by 10° interval in dots on x/L_max_ plot, WT in black, Cut remained in red. **(f)** Psmad signal of each embryos averaged by 10° interval in dots on x/L plot, WT in black, Cut remained in red. Each dot corresponds to an average of Psmad signals of all cells within every 10° along the margin within one embryo. **(g)** – **(i)**: Fitted curves and point-wise SEs for population mean morphogen gradients of Cut and WT on *x/L* plot and x/*L*_*max*_ plot. **(g)** SE of Cut and WT population mean morphogen gradients on x/L_max_ plot (blue) vs on x/L plot (magenta). **(h)** Curve fitting of population Psmad gradients on x/L_max_ plot. Average Psmad signals of every 10° along the margin of each WT embryo are fitted with a smooth curve using the Hill equation and plotted in black line on x/L_max_ plot. **(i)** Curve fitting of population Psmad gradients on x/L plot. Average Psmad signals of every 10° along the margin of each WT embryo are fitted with a smooth curve using the Hill equation and plotted in black line on x/L plot. **(j)** – **(l)**: Ratio of curve-fit parameters vs L along the Cut and WT embryo margins. **(j)** Scatter plot of the curve-fit parameter a vs L fitting with a line in blue, used to calculate the scaling power of parameter a. **(k)** Scatter plot of the curve-fit parameter b vs L fitting with a line in blue, used to calculate the scaling power of parameter b. **(l)** Scatter plot of the curve-fit parameter c vs L fitting with a line in blue, used to calculate the scaling power of parameter c.

According to Figure 1, the distribution of phenotypes shows that scaling is not guaranteed, at least for the distribution of sizes within the whole population. Similarly, the BMP signaling pattern demonstrates increased embryo-embryo variability after yolk removal. Due to this, we measured scaling properties from two populations of embryos- the complete population- including those without appreciable signaling, and a population filtered to only include embryos that exhibit measurable Psmad signaling. The rules for the filtered population are as follows:

1. The Cut embryos with peak Psmad relative intensity beyond the range of the peak Psmad labeling intensity of WT embryos ([0.60, 1.43] after normalization) are discarded.
2. If the difference of Psmad relative intensity does not exhibit a sufficient gradient between ventral and dorsal half is less than 0.2 after normalization, the Cut embryo is discarded. As a result, in the filtered data-set we retained 20 out of 50 Cut embryos.

Figure 2b shows the relative size of fluorescently stained WT and Cut embryos. Most of the Cut embryos are 20~30% smaller than WT embryos in length, and 50~70% smaller in volume. The size variation within WT embryos is about 20%. The Cut embryos are 20% smaller than WT on average and the smallest Cut embryo is 50% smaller than WT. Sample size of WT and Cut are 24 and 50 respectively.

#### Metric 1: Morphogen gradients on relative and absolute position plots

Figure 2c-f plots Psmad gradients of WT and Cut embryos after the sorting out non-signaling. Compared to Psmad gradients of Cut and WT embryos on the absolute position plot (Figure 2c), the Psmad gradients of WT and Cut embryos converge better on an x/L plot (Figure 2d). The scatter plot of average Psmad relative fluorescence of each WT and Cut individual embryo is shown on an x/L plot. Figure 2f also shows a better overlap than the overlap on an x/*L*_*max*_ plot (Figure 2e), especially in the range of 0.2 to 0.4. Recall that *L*_*max*_ is the single biggest embryo length measured. A student’s t-test was used between WT and Cut Psmad relative fluorescence at each node for the 9 ventral lateral space nodes on the x/L plot and on the x/*L*_*max*_ plot. The p values of the t test for WT and Cut Psmad labeling intensities at all the 9 space nodes on the x/L plot are >0.05. This indicates that we could not reject the null hypothesis for a difference between Psmad of WT and Cut embryos when they are all plotted on a scaled x coordinate. In contrast, the p-value for the t-test is below 0.05 at 8 out of 9 space nodes on x/*L*_*max*_ plot, indicating the WT and Cut Psmad labeling intensities are significantly different at most of the ventral lateral regions on the x/*L*_*max*_ plot. This comparison confirms the better overlap of WT and Cut Psmad gradients on the x/L plot.

The improvement of overlap between the WT and Cut Psmad gradients on the x/L plot as opposed to the x/*L*_*max*_ plot is a hallmark of scaling. The Psmad gradients of each embryo and the population averages are fit to the Hill equation and quantitatively evaluated for scaling by comparing the morphogen gradient curves of WT and Cut embryos plotted in absolute position vs in relative position (Figure 2g - l).

#### Metric 2: Point-wise scaling error

Figure 2g shows that point-wise scaling error (SE) of fitted curves of population averages between WT and Cut embryos on an x/L plot is smaller than that on *x/L*_*max*_ plot (throughout lateral regions), indicating a better overlapping/convergence of the morphogen gradients of WT and Cut on x/L plot over x/L_max_ plot. Note, SEs at dorsal and ventral ends are not shown because the curve at these two ends are very flat.

#### Metric 3: T test of curve parameters

We then compared the shapes of gradient curves by the curve parameters of the fitted Psmad gradient curves of each embryo. As shown by t test results in Table 1, all three parameters of the function that fit morphogen gradients of WT and Cut embryos on x/L plot do not significantly differ. This indicates that the shape and amplitude of morphogen gradient curves of WT and Cut embryos do not differ. This is consistent with Pearson’s correlation test results for curve-fit parameters in Table 2 and the slope of curve parameters vs L in Figure 2j - l, that curve-fit parameters do not correlates with size significantly, indicating that the shape and amplitude of curves that fit morphogen gradients on the x/L plot do not significantly change with egg size.

**Table 1.** t test of curve-fit parameters of Cut vs WT on x/L plot.

|  | p value |
| --- | --- |
| $a_{WT} \text{ vs } a_{Cut}$ | 0.820 |
| $b_{WT} \text{ vs } b_{Cut}$ | 0.30 |
| $c_{WT} \text{ vs } c_{Cut}$ | 0.44 |

**Table 2.** Correlation Test for curve-fit parameters vs L.

|  | Correlation Coefficient | p value |
| --- | --- | --- |
| a vs L | -0.18 | 0.24 |
| b vs L | -0.12 | 0.44 |
| c vs L | 0.29 | 0.059 |

The results from the 3 metrics and the normal phenotypes of Cut embryos indicates that the scaling of DV patterning can be traced back to the scaling of morphogen gradients in zebrafish.

### Interspecies scaling between zebrafish and giant danio embryonic DV patterning

Giant danio (Gd) and *Danio rerio* zebrafish are closely related species. They belong to in the same family of Cyprinidae and subfamily of Danioninae. Specifically, the zebrafish line we use to compare with Gd is Tuebingen (Tu). Gd eggs are, on average, 20% larger and 1.5~3 times larger as adults than zebrafish. As shown in Figure 3a and b, we observed that the egg size of giant danio is bigger than that of zebrafish. To determine the level of pattern likeness and scaling, we measured Psmad morphogen gradients of both and quantified the degree of scaling of their morphogen gradients using the metrics described above.

**Figure 3.**
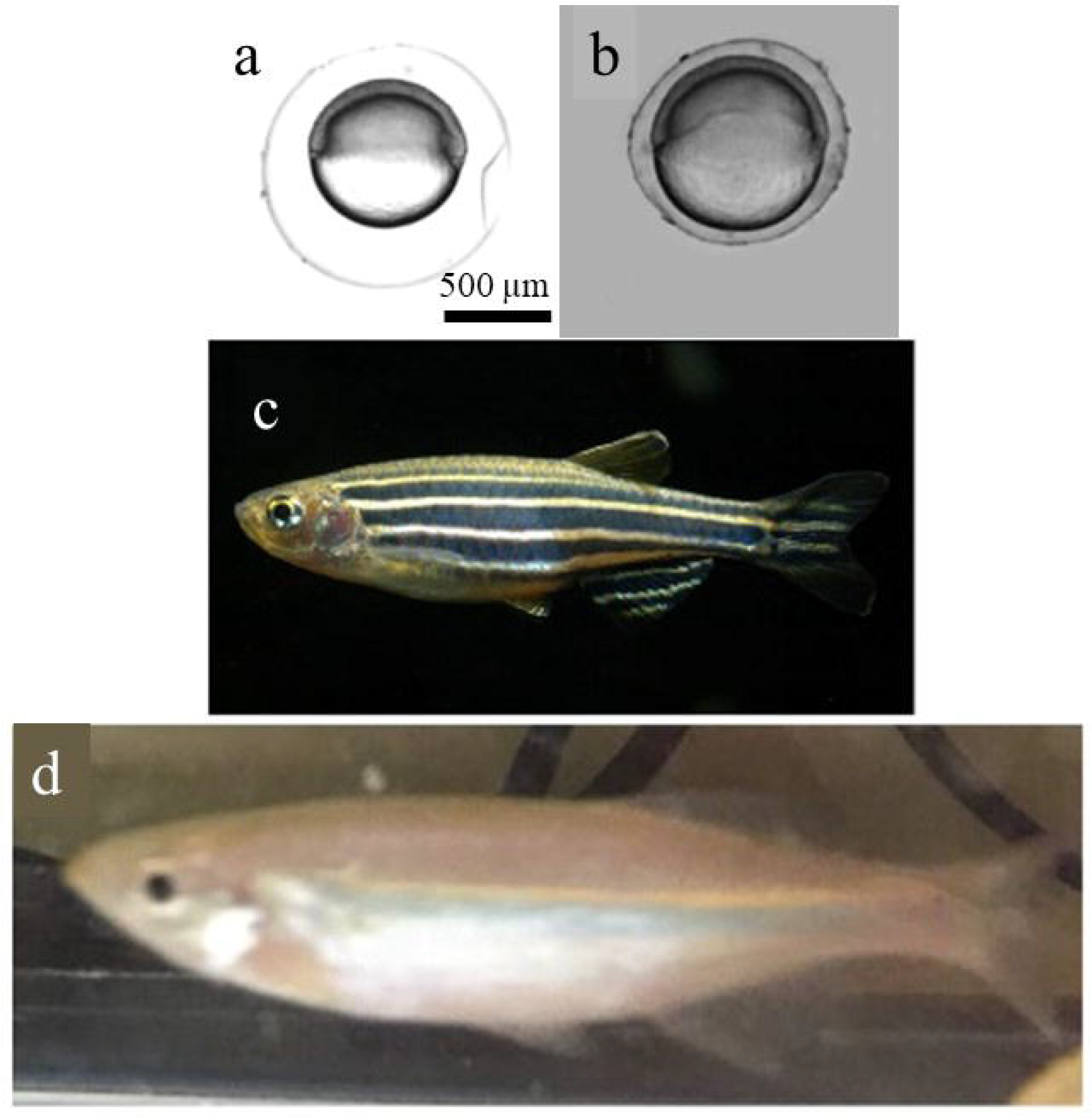
The Gd embryos are significantly bigger than Tu. **(a)** Bright-field image of one Tu embryo at shield stage. **(b)** Bright-field image of one Gd embryo at shield stage. **(c)** Bright-field image of one Tu embryo at 1 dpf. **(d)** Bright-field image of one Gd embryo at 1 dpf.

#### Metric 1: Morphogen gradients on relative and absolute position plots

We quantified the Psmad gradient from fluorescence images of the Psmad stained Tu and Gd embryos. Figure 4a shows an example point cloud of Tu and Gd embryos with Psmad labeling intensity in color. We measured Psmad morphogen gradients of 55 Tu and 15 Gd embryos and the average size of Gd is about 24% larger than that of Tu (Figure 4b). The Psmad gradients of Tu and Gd embryos exhibit less overlap on the x/L plot (Figure 4d) than they do on the x/*L*_*max*_ plot (Figure 4c). This indicates that morphogen gradients of Tu and Gd do not scale. To further evaluate scaling, the Psmad gradients were fit to the same function as before and compared.

**Figure 4.**
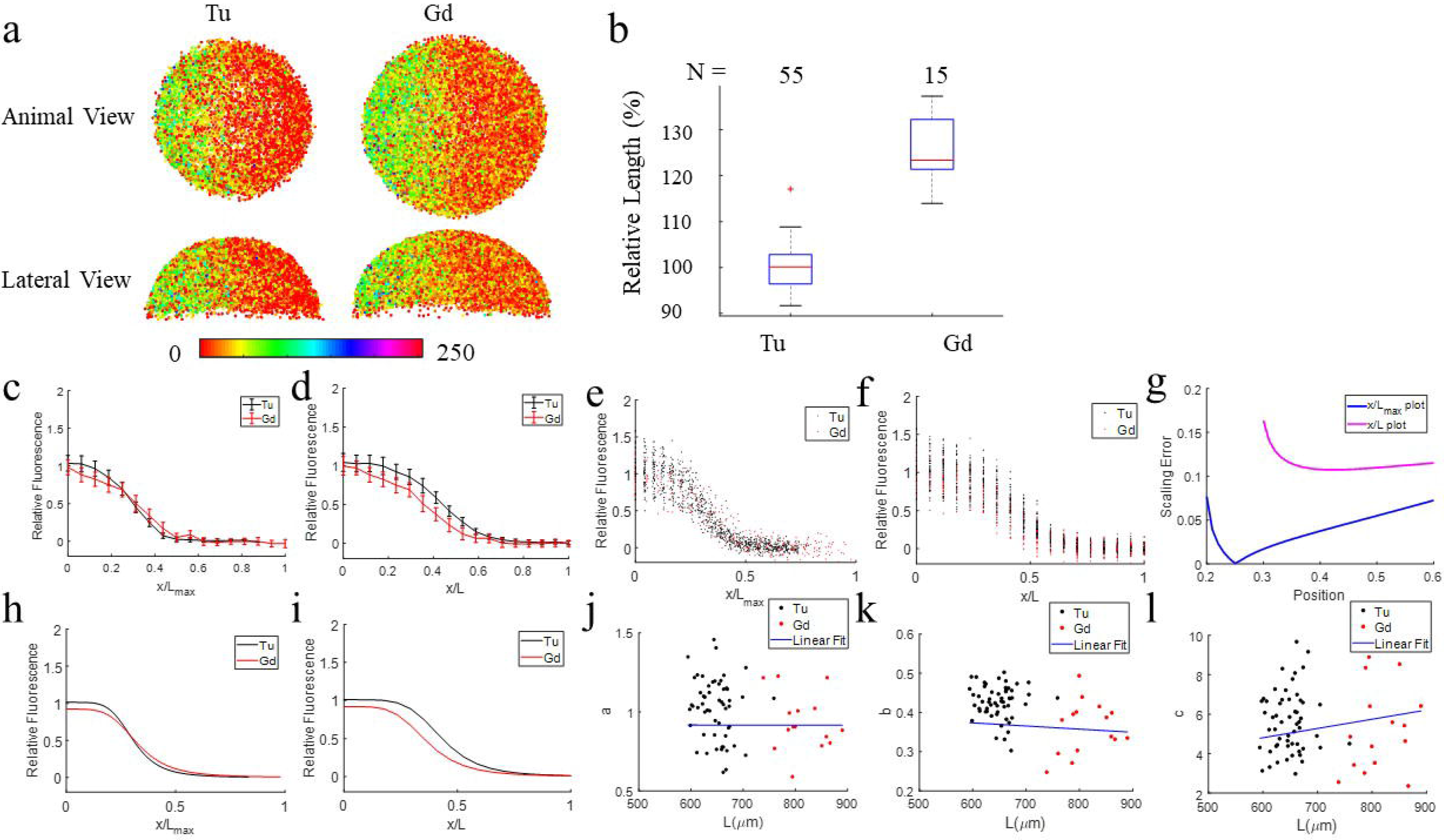
The Psmad gradient profiles of Gd and Tu do not scale. **(a)** The animal and lateral views of Psmad stained 6 hpf embryo point clouds of Tu and Gd after image processing with Psmad labeling intensity in color. **(b)** The distribution of the relative vegetal margin radius length of Tu and Gd. **(c)** – **(e)**: Psmad gradient profiles of Gd (red) vs Tu (black) embryos. **(c)** Population mean Psmad gradients of Gd and Tu on x/L_max_ plot with variation shown by error bars. In each embryo, the Psmad levels of cells are averaged among cells that are nearby within an interval of 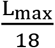 in length. Psmad labeling intensities of all embryos of the same group at 18 the same absolute position are then averaged to plot the errorbar on each position node. **(d)** Population mean Psmad gradients of Gd and Tu on x/L plot with variation shown by error bars. **(e)** Psmad signal of each embryos grouped by 10° interval in dots on x/L_max_ plot, Tu in black, Gd in red. **(f)** Psmad signal of each embryos averaged by 10° interval in dots on x/L plot, Tu in black, Gd in red. Each dot corresponds to an average of Psmad signals of all cells within every 10° along the margin within one embryo. **(g)** - **(i)**: point-wise SEs for population mean morphogen gradients of Gd and Tu after curve fitting on x/L plot and x/L_max_ plot. **(g)** SE of Gd and Tu population mean morphogen gradients on x/L_max_ plot (blue) vs on x/L plot (magenta). **(h)** Average Psmad signals of every 10° along the margin of each Tu embryo are fitted with a smooth curve using the Hill equation and plotted in black line on x/L_max_ plot. The smooth curve that fits the individual average Psmad signals of Gd embryos are plotted in red line. **(i)** Curve fitting of population Psmad gradients on x/L plot. **(j)** – **(l)**: Ratio of curve-fit parameters vs L along the Gd and Tu embryo margins. **(j)** Scatter plot of the curve-fit parameter a vs L fitting with a line in blue, used to calculate the scaling power of parameter *a*. **(k)** Scatter plot of the curve-fit parameter b vs L fitting with a line in blue, used to calculate the scaling power of parameter b. **(l)** Scatter plot of the curve-fit parameter c vs L fitting with a line in blue, used to calculate the scaling power of parameter c.

#### Metric 2: Point-wise scaling error (SE)

Figure 4g shows that point-wise SE of fitted curves of population average between Tu and Gd on x/L plot is bigger than that on x/*L*_*max*_ plot (throughout lateral regions), consistently indicates less overlapping of the morphogen gradients of Tu and Gd on the x/L plot as opposed to the x/*L*_*max*_ plot.

#### Metric 3: T test of curve parameters

We also compared the shapes of gradient curves by the curve parameters, the amplitude parameter a, and the slope parameters b and c. In Table 3, the p value of b is less than 0.05. To achieve power as high as 0.8 for the t test of parameter b given the observed mean and variance of data, the sample size for each group should be at least 9. Here, sample size of Tu and Gd embryos are 55 and 15 respectively, both are higher than 9. Power analysis is conducted for the t test of parameter b given sample size as 15. The observed power for the t test of parameter b is 0.95, very high. This means that the parameter corresponding to position shift of the curve, b, of Gd and Tu morphpogen gradients on the x/L plot is significantly different by t-test, indicating that the shapes of morphogen gradients of Tu and Gd are significantly different and that they do not scale. This is consistent with the Pearson’s correlation test results for the curve-fit parameters in Table 4 and the slope of the curve-fitting parameters vs L in Figure 4j - l.

**Table 3.**
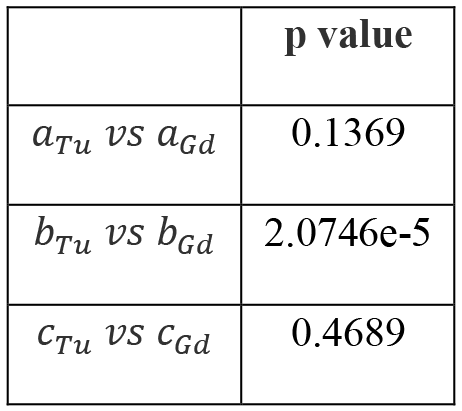
t test of curve-fit parameters of Tu vs Gd for x/L plot.

|  | p value |
| --- | --- |
| $a_{Tu} \text{ vs } a_{Gd}$ | 0.1369 |
| $b_{Tu} \text{ vs } b_{Gd}$ | 2.0746e-5 |
| $c_{Tu} \text{ vs } c_{Gd}$ | 0.4689 |

**Table 4.**
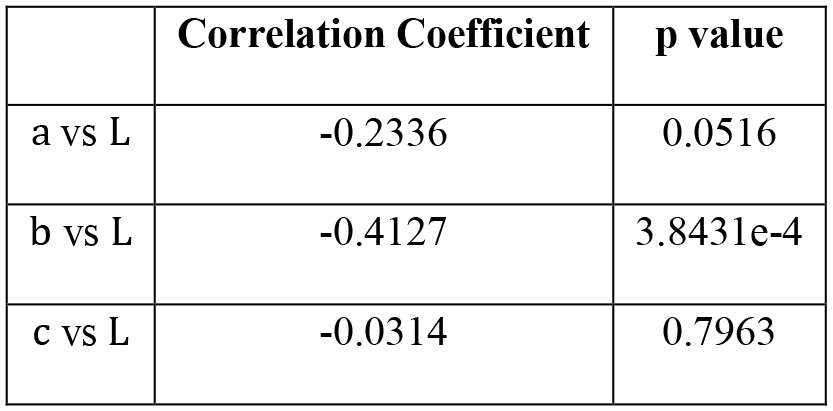
Correlation Test for curve-fit parameters vs L.

The above metrics consistently demonstrate that the DV patterning and corresponding morphogen gradients of Gd embryos do not appear to scale with zebrafish embryos.

## Discussion

The experimental data for Cut and WT zebrafish embryo images and quantitative analysis of the morphogen gradients leads to the conclusion that zebrafish embryos possess some degree of scaling with egg size. This data is consistent with the observed phenotypes- and the distribution of disrupted patterns is consistent with distribution of phenotypes. The analysis herein does not identify the mechanisms that lead to scale invariance, however future studies that couple these imaging approaches, genetic perturbations of BMP pathway components and mathematical modeling will be used to identify these mechanisms in the future.

In contrast to the results of embryonic scaling in surgically perturbed populations of *Danio rerio*, the Psmad gradient does not appear to scale between closely related species. Gd embryos are significantly bigger than the Tu line of *Danio rerio* and the morphogen gradients do not scale. The gradients exhibit greater overlap when plotted on absolute coordinates or when each is normalized to the maximum sized individual within each population (x/*L*_*max*_).

The next challenge in the study of scale invariance is to move from describing the patterns of scaling relationships to understanding the underlying processes that create the patterns. In the future, possible mechanisms of intraspecies scaling can be investigated. The fact that BMP gradients in Gd and Tu do not scale may have to do with the time it takes for BMP gradient formation and morphogen spread. The time-scale for molecular diffusion is proportional to 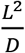 (where *D* is the diffusion coefficient) [**Error! Bookmark not defined**.]. We know that the average circumference length the of Gd embryo is about 20% longer than that of Tu. So, if the values of diffusion coefficient are about the same in Gd and Tu, the time for BMP to diffuse across the field of cells increases by about 44% on average. Furthermore, if the decay rates and production rates of extracellular regulators are about the same in Gd and Tu, the BMP gradient will be shorter as more ligand will decay before it has time to diffuse the greater distances around the embryo.

In summary, this work describes our methodological advancements in the study of scaling in zebrafish embryos, and communicates our insights on DV pattern scale invariance within and between fish species. We conclude that DV pattern scale invariance in zebrafish embryos can be traced back to the intraspecies scaling of BMP gradients, and there is no scaling of BMP gradients between zebrafish and giant danio.

## Materials and methods

### Embryonic surgery and confocal fluorescence imaging

#### Vegetal yolk removal surgery

Embryo size is reduced by removing vegetal yolk during the 8-cell to 256-cell cleavage stage at room temperature. Maternally provided dorsal determinants are known to activate zygotic gene expression cascades required for the formation of the dorsal organizer [28, 29, 30, 31]. At 20 minute-post-fertilization (mpf), a microtubule array forms and Syntabulin transports the dorsal determinants to the plus end of microtubules. Around the 2-cell stage (60 mpf), Syntabulin is not detectable by immunostaining and whole embryo lysate immunoprecipitation, indicating the degradation of Syntabulin and the release of the dorsal determinants into cells that are eventually incorporated by dorsal blastomeres [32]. Therefore, the surgery is conducted after the 4-cell stage in order to not disrupt the delivery of dorsal determinants from yolk to cells. Chorion protects the embryo from being destroyed by surface tensions and is not removed during the yolk removal surgery. To do the surgery, we hold a tungsten needle (0.125 mm ultra-fine) and pierce across the chorion without destroying it while stabilizing the embryo by holding another part of the chorion with forceps. Then we poke a hole on the vegetal yolk with the needle. The surgery is conducted in 1X E3 medium in petri dishes bedded with 1% agar. The egg size is controlled by removing various amounts of yolk. Yolk removed embryos are called “Cut” while controls that do not have any treatment are called wild-type (WT).

#### Phenotype observation

Embryos grow in an incubator at 28 °C for 24 hours after the surgery. In the 1X E3 solution, 10U/ml Penicillin and Streptomycin are added to protect embryos from bacterial infection and Methylene Blue is added to prevent from fungus growth. Images of the embryos are taken under bright-field microscope at shield stage and 1 day-post-fertilization (dpf) to observe the morphological phenotype along the DV axis.

#### Imaging of Psmad1/5

Embryos are fixed at shield stage with 4% paraformaldehyde at 4 °C, blocked in NCS-PBST (10% fetal bovine serum, 1% DMSO, 0.1% Tween 20 in PBS), and probed overnight with a 1:100 dilution of anti-phosphoSmad1/5/9 antibody (Cell Signaling Technology, #13820s), followed by a 1:500 dilution of goat anti-rabbit Alexa Fluor 6470 conjugated antibody (Thermo Fisher Scientific, Rockford, IL; Cat# A-21244, RRID: AB_2535812). Nuclei are stained by sytox orange. Embryos were mounted in BABB (benzyl alcohol (Sigma B-1042) and benzyl benzoate (Sigma B-6630), 1:2 ratio) and scanned using a Zeiss LSM 800 confocal microscope with a 20X water lens. The detailed protocol of immunostaining and imaging has been published [27]. Here, Psmad fluorescence intensity is used as a readout of extracellular BMP in zebrafish.

### Quantification of morphogen gradients

#### Computational image analysis

We quantified the Psmad gradient through an image analysis pipeline that includes cell segmentation, preprocessing and image registration, and data extraction along the embryo margin [26]. The marginal circumference of the embryo image is measured by extracting a margin region with a thickness of 60 μm and averaging the circumference of the shape. The measurement of embryo size here is half of the margin circumference, denoted as L. Note, *L*_*max*_ is the half margin circumference of the largest embryos. The circumferential position is normalized to *L*_*max*_. It is in the same range as the x-axis of a relative position plot x/L, [0,1], to compare quantities calculated from these two plots.

#### Data normalization and data reduction

Psmad gradient normalization for populations of images taken on different days are conducted using WT controls [27, 33]. Psmad gradients of Gd embryos are normalized within the population of Gd and not normalized against *Danio rerio* due to possible differences in antibody binding efficiency. Maxima of population averages for Danio rerio and Gd are both set to 1.

Some embryos are too severely damaged by the tungsten needle and do not progress through development. Embryos that are too damaged or exhibit no Psmad signaling due to development arrest are eliminated from the population for quantification of Psmad signaling.

#### Curve fitting morphogen gradients

Multiple measures of scaling are used to compare the gradients - pointwise measurements for local intensity differences and parameters from functions that are fit to the data. Previous work for morphogen gradient quantification relied on comparison of the length scale of the gradient determined by fitting the data to an exponential function. These functions do not apply here because the gradient shape is not exponential. Many mathematical functions including polynomial, exponential, Fourier, Gaussian, etc. were tested and we found that a variant of the Hill equation fits the shape of morphogen gradients best (*R*^2^ > 0.95) and was selected to fit the data:

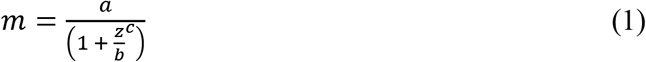

m is the concentration of morphogen and z is the position, either relative position 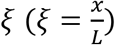 or the absolute position scaled to [0,1] by 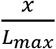. Parameter *a* represents the amplitude of the curve while the slope depends on the value of *b* and *c*. The curve has a fixed point at 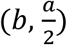.

### Ethical approval of using animals

The experiments were approved by Purdue Animal Care and Use Committee (PACUC) with Animal Use Qualification Form on file. We confirm that all experiments were performed in accordance with relevant guidelines and regulations.

### Data availability statement

All the 3D point cloud images of embryos and the data matlab files for the study of intraspecies and interspecies scaling are stored in a dropbox shared folder. The link to it is https://goo.gl/mcxY3R. It is also included in the supplemental files.

## Acknowledgements

I thank Dr. Pascal Lafontant’s group for providing me giant danio embryos. I acknowledge Dr. Mary Mullins, Francesca Tuazon and Dr. Joseph Zinski for help in developing the protocol for reducing the embryo size. I thank Dr. Bobby Madamanchi for offering feedback for the manuscript. We thank the NIH and Grant number HD073156 for support.

## Author contributions statement

Y.H. conceived the experiments, conducted the experiments and analyzed the results. D.U. also conceived the experiments and analyzed results. Y.H. and D.U. wrote the main manuscript text and prepared figures. All authors reviewed the manuscript.

## Additional information

Competing interests statement: the author(s) declare no competing interests.

